# High order structure in mouse courtship vocalizations

**DOI:** 10.1101/728477

**Authors:** Stav Hertz, Benjamin Weiner, Nisim Perets, Michael London

**Affiliations:** The Alexander Silberman Institute of Life Sciences, The Hebrew University of Jerusalem, Jerusalem, 91904 Israel; Edmond and Lily Safra Center for Brain Sciences, The Hebrew University of Jerusalem, Jerusalem, 91904 Israel

**Keywords:** mouse ultrasonic vocalization, information theory, classification, syntax

## Abstract

Many complex motor behaviors can be decomposed into sequences of simple individual elements. Mouse ultrasonic vocalizations (USVs) are naturally divided into distinct syllables and thus are useful for studying the neural control of complex sequences production. However, little is known about the rules governing their temporal order. We recorded USVs during male-female courtship (460,000 USVs grouped into 44,000 sequences) and classified them using three popular algorithms. Modeling the sequences as Markov processes revealed a significant temporal structure which was dependent on the specific classification algorithm. To quantify how syllable misclassification obscures the true underlying sequence structure, we used information theory. We developed the Syntax Information Score and ranked the syllable classifications of the three algorithms. Finally, we derived a novel algorithm (Syntax Information Maximization) that utilized sequence statistics to improve the classification of individual USVs with respect to the underlying sequence structure.

## Introduction

Mice emit ultrasonic vocalizations (USVs) in various behavioral contexts (Sewell, 1968; Sales, 1972; Holy and Guo, 2005; Portfors, 2007; Seagraves et al., 2016). Recently, this behavior has gained interest as a proxy model for speech and language (Fischer and Hammerschmidt, 2011; Arriaga et al., 2012; Portfors and Perkel, 2014; Fisher and Vernes, 2015; Castellucci et al., 2016) and as a tool for behavioral phenotyping of neurodevelopmental disorders (Scattoni et al., 2008; Yang et al., 2015; Egnor and Branson, 2016). However, to fully gain access to the advantages provided by the information hidden in USVs, we need adequate methods to analyze this complex signal.

Observing the spectrogram of the sound signal (Fig. 1A), it is easy to appreciate that it is composed of distinct individual syllables where the power in the ultrasonic range (> 20 kHz) is made of continuous stretches of positive power (USVs) or zero power (silence). The periods of silence between USVs (inter-syllable intervals, ISIs) follow a typical distribution with several distinct peaks, suggesting a prototypical process of producing these sounds. Thus, it is possible to parse the acoustic signal into individual USVs and USV sequences based on the distribution of ISIs (Fig. 1, and in: Holy and Guo, 2005; Chabout et al., 2015; Castellucci et al., 2016).

**FIGURE 1.**
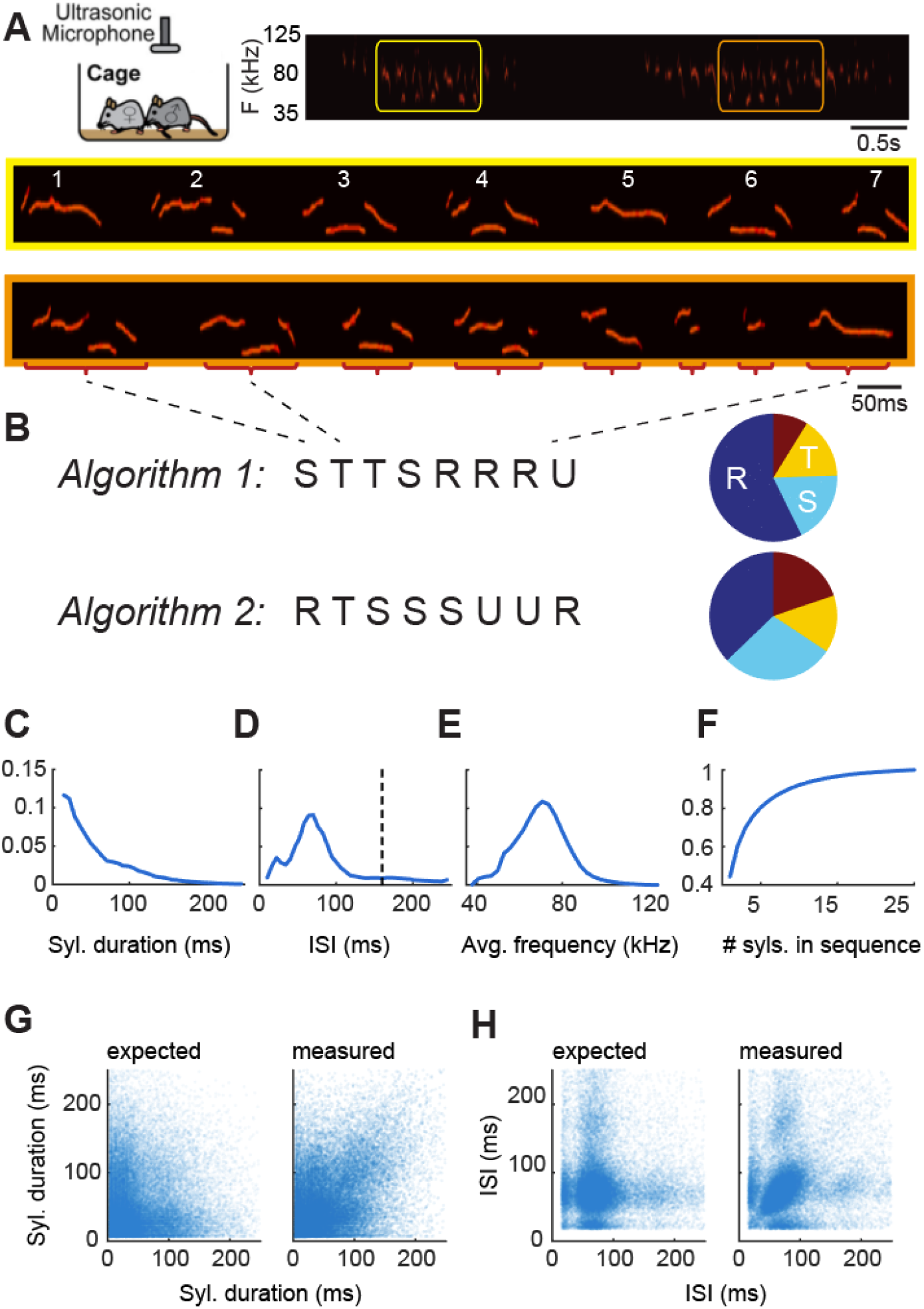
Parsing and classification of USV sequences. **A** Top: Example spectrogram of USV recording composed of two sequences. Bottom: example USV sequences composed of several syllables, displaying richness of shapes. Some shapes have distinctive features that may lead to natural categorization, such as sudden jumps in frequency (e.g. syl. #3, #6). However, syllables differ in many ways including the duration of sub-components (e.g. 3,6,7) or number of sub-components (e.g. 1,2 or 2,4 or 3,4). Other features, e.g., syllable duration, have a continuous distribution over a wide range and their use for distinction between classes is less obvious (see also panel **C**). **B** Classification examples by two illustrative algorithms. Syllables are classified into one of four classes, marked by the letters: ‘RSTU’. Algorithm 1 classifies by the number of sub-components while Algorithm 2 by syllable duration. The distributions of the classes, presented in the pie charts, indicates that the algorithms are not homologous. (**C-F**) Distributions of syllables parameters collected from ~500,000 syllables and ~50,000 sequence. **C** syllable duration, **D** inter-syllable interval (ISI), **E** average syllable frequency and **F** cumulative number of syllables in a sequence. **G** correlation pattern between duration of adjacent syllables in sequences (left: expected pattern based on **C**, right: actual pattern in data). **H** same as **G** but for ISIs.

A closer look at these individual vocalizations suggests the existence of classes. Many USVs have a rather simple form, composed mainly of narrow band frequency sweeps. Some other USVs are composed of substructures of various sweeps, each of which can change its length and main frequency, resulting in a wide spectrum of shapes (Fig. 1A). Therefore, while the process of parsing the sound into individual USVs is primarily one of overcoming technical obstacles, the classification of the individual USVs into syllable classes based on their acoustic features presents complex, fundamental challenges.

In the absence of “ground truth” (in sharp contrast to human speech) various approaches have been taken in developing methods of classification. For example, Holy and Guo (Holy and Guo, 2005) and later (Chabout et al., 2015, 2016) have used frequency jumps as the main feature of differentiating individual USVs, and have classified them according to the number and direction of the jumps while ignoring other features (e.g. duration). Alternatively, other algorithms have taken unsupervised learning approaches without deciding upfront on hardwired features (e.g. Burkett et al., 2015; Van Segbroeck et al., 2017). It is important to realize that when the same ensemble of USVs is classified by different algorithms there might not be a one-to-one mapping between the resulting labeling (Fig. 1B). This important issue will have significant consequences as discussed below.

Figure 1A highlights another property of mouse vocalization, the existence of complex syllable sequences. Vocal communication systems in other species (e.g. human speech, bird songs) are also based on sequences of sound units. Words are composed of syllables, and sentences are composed of words. Songbirds are known to produce diverse and complex sequences or “songs” (Doupe and Kuhl, 1999; Brainard and Doupe, 2002; Jarvis, 2004). Some of these songs contain hierarchical acoustical units: notes, syllables and motifs (Okubo et al., 2015) which can be very stereotypic (e.g. Zebra finch (Price, 1979; Williams and Staples, 1992) or have an underlying complex syntax (e.g. Bengalese finches (Hosino and Okanoya, 2000; Okanoya, 2004)). USVs produced by male mice during courtship share some of the syntax characteristics of songbirds (Holy and Guo, 2005; Sugimoto et al., 2011; Chabout et al., 2015).

Taken together, the non-homologous assignment of labels by different classification methods and the existence of complex structures in USVs sequences, may lead to very different statistical properties of labeled sequences from the same data. For example, the distribution of the number of syllables in each class could take many forms and similarly, the probability of a syllable pair appearing together in a sequence may vary. Therefore, the selection of the classification algorithm may lead to different scientific conclusions and interpretations.

Here, we show that this undesirable consequence of having different syntaxes from the same data, could, in fact be useful in selecting and improving classification methods. This is based on the observation that a syntax imposed by an algorithm dictates how well it predicts the label of the next syllable in a sequence. A “meaningful” classification should have a high predictive power, which could then be taken as a measure for the classification quality. Moreover, the information that is present in the sequence structure could improve the classification. Using information theory tools, we examine these ideas by comparing the predictive power of various classification methods and we suggest a simple way to incorporate optimization of the predictive power as an integral driving force of a classification algorithm.

## Results

### Analysis of basic USV properties suggests high-order structures

To study the differences between classification algorithms we compiled a database from our USV recordings. The recordings were made during sessions of interaction between adult male and female mice for a total of 113 hours. We developed an analysis toolkit (written in Python; available online) with a parsing algorithm to detect in the audio files the exact start and end times of each USV and each sequence of USVs. Applying this algorithm to our recordings, we extracted 461,927 USVs, which were then stored in the database along with their features. The individual USVs where grouped into 44,358 sequences.

Figures 1C-F show the analysis of the basic properties of USVs in our database. The three main properties that we focused on were: syllable duration, inter-syllable interval (ISI) and syllable mean frequency. For each one of these properties we show the general distribution and the correlations. The occurrence frequency of syllable duration fits a monotonically decreasing exponential (adjusted R-square: 0.9934) (Fig. 1C). The distribution of ISIs had two peaks in agreement with previous reports (Chabout et al., 2015; Castellucci et al., 2016), however, in our hands the two peaks occurred at shorter durations than previously reported. The first peak was at 20ms and a larger peak at around 70ms. Careful observation of the ISI distribution of different individual mice revealed a more complex picture, in which some mice had these double peak distributions while others did not (Fig. S1).

We found significant correlation in the distribution of syllable duration (r = .43, p < .001), such that short syllables tend to follow short syllables and long ones to follow long ones (Fig. 1G). Similar results were found for the correlation of ISIs (Fig. 1H, r = .16, p < .001). In conclusion, the existence of correlations already at this level of analysis (i.e. before classification) suggests that USVs are not emitted independently of each other, and that USV sequences have a non-obvious temporal structure.

### Classification of the same USVs with different algorithms

In order to test if different classification methods are homologous, we tested three classification algorithms that were recently published. The methods that we chose were: MSA v1.3 (Chabout et al., 2015), VoICE (Burkett et al., 2015) and MUPET (Van Segbroeck et al., 2017) (Fig. 2). We chose these algorithms because of the following reasons: (1) they represent different approaches to classification (2) they require relatively low manual involvement, and (3) the published algorithm provided code that could be applied to our database with relatively minor modifications (see Materials and Methods). Here we refer to them as iMSA, iVoICE and iMUPET to emphasize that we used the modified algorithm, which was inspired by the original algorithm.

**FIGURE 2.**
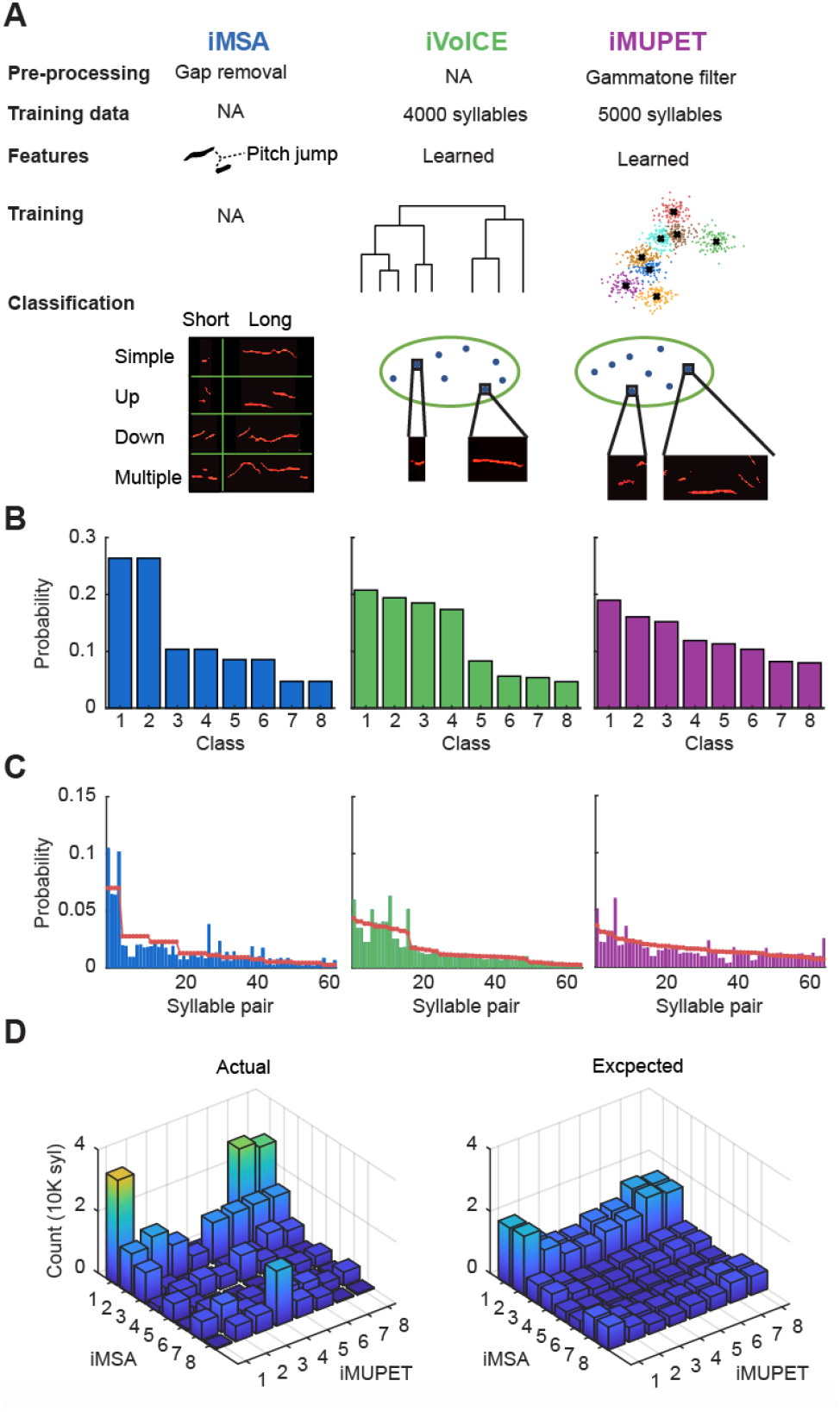
Classification of the same data with different algorithms. **A** A concise representation of the different algorithms that were adapted for this study. iMSA (left column) first pre-processes the data for gap removal. It requires no training data and classifies the syllables into classes based on their pitch jumps. The four basic pitch jump classes: Simple (no pitch jump), Up, Down and Multiple. Each is then divided into two according to its median syllable duration for a total of eight classes. iVoICE performs hierarchical clustering on a training subset of 4000 syllables resulting in eight centroids that are then used to classify the rest of the syllables based on a similarity measure. iMUPET algorithm performs a preprocessing gammatone filter on all syllables, and then uses the k-means algorithm to create centroids from 5000 syllables. These centroids are used to classify the rest of the syllables based on the cosine distance between the filtered syllable and the centroid. **B** The distributions of the classified USVs from the database are shown for each algorithm, iMSA produces the most nonuniform distribution and iMUPET the most uniform one. The difference between the distributions means that the algorithms are non-homologous. **C** The distributions for pairs of USVs are shown for each algorithm. The red line depicts the predicted distributions derived from **B** assuming independence. The histograms are sorted by the expected distribution. **D** Joint distribution of the classifications of iMSA and iMUPET for all USVs in the dataset (left) and the expected joint distribution assuming independence (right).

Figure 2A describes the key properties and the workflow of the three methods. In short, iMSA is based on hardwired features (pitch jumps), while iVoICE and iMUPET apply unsupervised methods to “learn” the key features. This learning is done on a learning set that is limited in size to a few thousands of USVs, resulting in a set of centroids which represent the different classes. iVoICE uses an hierarchical clustering strategy, and iMUPET uses the k-means clustering algorithm (MacQueen, 1967) with a given number of clusters. The classification of a USV to a class is obtained by assigning each observed USV to the closest class representative (centroid) using some metrics (spectral similarity in the case of iVoICE and cosine distance in iMUPET). Two example centroids are shown for each of the algorithms.

To compare the algorithms, we ran all of them with eight classes. This gave a good balance of rich classification on the one hand, while still enabling the collection of enough higher order statistics, which will be important for later analysis. For the iVoICE and iMUPET algorithms, the number of classes is a natural parameter. However, iMSA assumes only four classes, so we obtained eight classes by splitting each class into two according to the median syllable duration (Materials and Methods).

Figure 2B shows the distribution of the number of syllables assigned to each of the eight classes for the three algorithms. Clearly, iMSA produced a distribution that is less uniform than the other two methods (iVoICE, iMUPET). For iMSA, the proportion of USVs with no pitch jump is somewhat larger than that of those with pitch jumps. Because iMSA classifies all syllables with no pitch jump as Simple (long or short), the first two classes occupy over 50% of the data and create a very non-uniform distribution. The difference between the distributions clearly shows that there is no one-to-one mapping between them (i.e. it is not that the class labels can be permuted to obtain the same classification) as suggested in the introduction.

Similar to the analysis of the basic properties, we also computed higher order distributions for each of the classifications. Figure 2C shows the distributions of pairs of syllables imposed by the three algorithms. In addition, the red line (which was used for sorting the histograms) represents the expected distribution derived from the distributions in figure 2B assuming independence. It is easy to see that in all cases there are large deviations from that expected distribution, implying that all algorithms capture some higher order structure of USV sequences. However, it is less obvious to deduce from that which algorithm captures more of this complexity.

The distributions in figure 2B clearly show that the different algorithms classify USVs differently and are not homologous. However, it is possible that they agree on the majority of the USVs and there is a relatively small group of USVs that are classified differently. To better test this possibility, we constructed the joint distribution between two of the algorithms. For each USV in the database we looked at the pair of its label, assigned by iMSA and iMUPET, and counted USVs for each of these pairs. If this option was true, one would expect that in each row in the joint matrix there would be one column with significantly high count, however as depicted in Figure 2D this is not the case (except from the pair of class 8 of iMSA and class 4 of iMUPET). For most classes of iMSA there is a fairly uniform distribution of the count over the iMUPET classes and vice versa. This distribution further strengthens the conclusion that the algorithms are not homologous.

### Predictive power of classification algorithms

Figure 2C suggests that the difference between the distributions imposed by the algorithms on the same data may be used to quantify the differences between them. Based on this observation we propose a framework for evaluating classification algorithms. The guiding principle is that a classification that exposes regularities in vocalization sequences is more likely to capture their underlying statistical structure. The better the statistical model of the USV sequences, the better the prediction it allows to draw about the future of the sequence. Therefore, we evaluate an algorithm by quantifying how well the syntax model it imposes predicts the future of the sequence.

Given a USV dataset and a classification method, the quantification process is done in two steps: (1) generating the syntax model and (2) calculating the model’s predictability. For step 1 (syntax model), we applied a classification algorithm to the USVs in the database and received sequences of class labels (i.e. sequences of labelled USVs). The transition probabilities of the labels were then modelled using an order-m Markov model (Fig 3A-C). This model describes the probability of a label in the sequence given the *m* preceding labels (i.e. the suffix). We represented the order-m Markov model as a Suffix tree of depth m. The probabilities of the next syllable given a suffix is stored in the leaves (Fig. 3; note in this notation a suffix is read backwards in time, from right to left).

**FIGURE 3.**
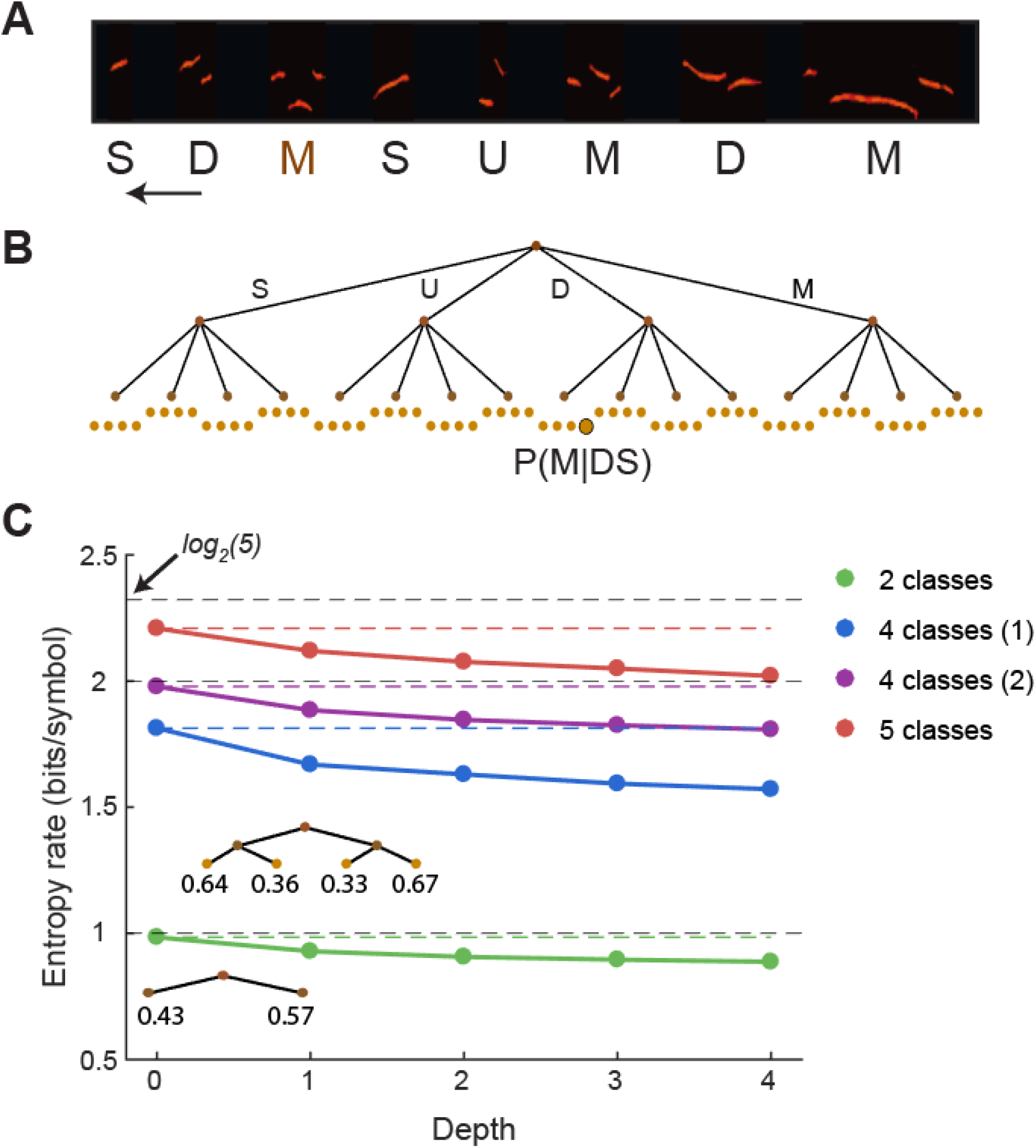
Computing the entropy rate of a classification method. **A** The classification algorithm is applied on every given sequence (in this example iMSA with four classes). **B** A suffix tree is constructed to hold the counts for a label given the suffix preceding it. Here an example for tree of depth D=2, the label M is preceded by the suffix DS (read from right to left) and thus the leaf M|DS counter is increased by 1 and the probability P(M|DS) is updated. This constructs a distribution from which the entropy rate of the syntax model is calculated. **C** Entropy rate values of four different classification algorithms (each marked by a different color; see text). The X axis represents the depth of the tree, the solid lines represent the entropy rate in each depth and the dotted colored lines represent the entropy rate of the 0^th^-order model. The dotted black lines mark the upper bound on the entropy rate for a given number of classes K (calculated as log_2_(K)). Inset trees represent the suffix trees with depth 1 and 2 for the algorithm that has two classes (green).

In step 2, for a given suffix tree we evaluate its predictive power by calculating the entropy rate of the order-m Markov model given by *H*(*Y*) = − ∑_*ij*_*μ*_*i*_*P*_*ij*_*logP*_*ij*_ where *μ*_*i*_ represents the probability of the *i*_*th*_ leaf and *P*_*ij*_ represents the probability of the *j*_*th*_ symbol in the *i*_*th*_ leaf. Roughly speaking, the entropy rate quantifies the amount of uncertainty regarding the class of the next USV given the syntax model and the labelled syllables in the suffix of length m. A low entropy rate (bounded by 0 bits/symbol from below) means a high amount of predictability while a high entropy rate means a high degree of uncertainty (and is bounded from above by the log of the number of possible labels).

### Definition of the syntax information score

Figure 2D demonstrates the application of this strategy for a few illustrative cases. We classified our USV database using four classification algorithms, and for each algorithm computed the entropy rate for Markov models of different suffix tree depths. The first algorithm (depicted in green) uses two classes: USVs containing pitch jump (J) or no pitch jump (N). The entropy rate of this model is bounded from above by 1 bit/symbol (dashed blue curve; when N and J appear independently and with equal probability). The distribution of the J and N syllables in our data was (43%,57%) and therefore with the 0^th^-order Markov model the entropy rate is 0.98 bits/symbol (Fig. 3C). For a 1^st^-order Markov model the entropy rate decreases to 0.93 bit/symbol due to the apparent tendency of J syllables to follow J (64%) and N to follow N syllables (67%). Computing the entropy rate for higher order Markov models shows modest decrease with the order, which saturates at order 4 (i.e. the additional contribution of the syllables beyond the first 4 in the suffix is negligible for predicting the next syllable).

Next, we considered two additional classification algorithms. The first is iMSA with the four original classes (Simple, Up, Down, and Multiple; depicted in blue). The second was a variant where the Up and Down classes were merged, and the Simple class was split into Short and Long syllables (depicted in purple). With four classes, the entropy rate is upper-bounded by 2 bits/symbol and for the 0^th^-order Markov model both schemes are slightly below that bound, because the distribution of labels is not uniform. Moreover, their 0^th^-order estimate of the entropy rate is not equal. Like in the previous case, the entropy rate of both algorithms decreases with increasing order of the Markov model. Note that in none of the cases does the entropy rate go to 0, meaning even given the full history of the sequence there is a significant randomness of the next syllable. Lastly, it is worth noting (and will be discussed further below) that the amount of drop in the entropy rate between the 0^th^-order model (1.81 and 1.98 respectively) and the 4^th^-order model (1.57 and 1.8 respectively) is not identical. Hence, the information gain between the case where the previous syllable is unknown and the case where we know the previous 4 syllables is not the same for the two models. We also considered an algorithm with 5 classes (Simple-Short, Simple-Long, Up, Down, Multiple; depicted in red), which strengthens our conclusions from the previous models, namely that the 0^th^-order entropy rate is close to log2(5) bits/symbol and that the reduction in entropy rate saturates at order-4 Markov model.

The analysis presented in figure 3C shows that entropy rate could be useful for comparing various classification algorithms. It also, however, highlights several subtleties. Firstly, the more classes the classification algorithm provides, the more likely that the entropy rate associated with it will increase. This is very natural, because the upper bound on the entropy increases as log_2_(number of classes) representing the fact that as the number of possible classes increase, so does the uncertainty regarding the class of the upcoming syllable. A meaningful comparison using entropy rate is possible only if the two algorithms use the same number of classes (or, alternatively, one could normalize by the upper bound).

The other important property of the entropy rate that is highlighted in the example, is that an algorithm can achieve a high predictability (low entropy rate) simply by assigning the same label to every USV independent of any feature. In such a case we know with certainty what will be the class of the upcoming syllable because it is always the same one. Unfortunately, this is the exact opposite result of finding regularities in the data. Rather, it is imposing ‘fake’ regularities by the algorithm. We conclude that merely using entropy rate as a measure for comparing algorithms is not ideal because the more non-uniform the distribution of classes at the 0^th^-order is, the lower the entropy rate, and the higher the predictability, irrespective of the true complexity of the sequences.

To overcome this problem, we note that the entropy rate of the 0^th^-order distribution is an inherent property of the classification algorithm. Since classification algorithms consider one USV at a time, it is unlikely that they introduce regularities of high order beyond those they impose on the 0^th^-order distribution. Therefore, a measure that is insensitive to the 0^th^-order distribution is more suitable for our purposes. We suggest using the KL divergence between the m^th^-order Markov model and the 0^th^-order (Kullback and Leibler, 1951; Cover and Thomas, 2005). This measure represents the information gain when changing our model from a 0^th^-order model to a higher order one. The KL divergence is calculated by 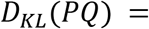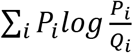 where in our case P is the higher order distribution and Q is the 0^th^-order distribution. The key idea here is that the KL divergence is bounded from above by the entropy of Q. Consider the case of two classification algorithms which use the same number of classes, where algorithm 1 imposes a very biased classification (tending to classify almost all USVs as the same class) and algorithm 2 results with a more balanced classification at 0^th^-order. The entropy rate of the 0^th^-order for algorithm 1 will be smaller than that of algorithm 2, which sets an upwards bound on the KL-divergence as described above. Therefore, algorithm 2 has a larger range to find regularities in the higher order distribution while algorithm 1 is penalized for the highly non-uniform distribution it imposes on the 0^th^-order distribution. Indeed, at the limit, an algorithm that assigns the same class to all USVs will have 0 entropy for the 0^th^-order distribution, enforcing KL divergence of 0 bits/symbol for any higher order model, and consequently the lowest predictability measure possible, in line with what we expect from our measure. We therefore suggest that the KL-divergence can serve as a good candidate to measure the predictive power of various classification algorithms. From here on, we will denote the Syntax Information Score (SIS; evaluated as *D*_*KL*_(*PQ*) from above, measured in units of bits/symbol) as our measure for comparing classification methods.

### Comparing the syntax information score for existing classification methods

We measured the SIS for the three classification algorithms presented in figure 2 applied to the USV database with eight classes. Figure 4A plots the entropy rates for the three algorithms for different depth of the suffix tree. Note that iMSA has the lowest 0^th^-order entropy as expected from Fig. 2B, while iMUPET, which has the most uniform distribution at 0^th^-order, yields an entropy rate of 2.9 bits/symbol (close to the maximum of 3 bits/symbol for 8 classes). However, it is easy to see the iMSA has the largest drop between the entropy rate at the 0^th^-order and the higher order ones. Fig. 4B shows the SIS for all three algorithms for D=1 and D=2. The graph shows that iMSA yields the highest SIS for both depths, despite its lower 0^th^-order entropy rate. iMUPET produces the second-best result. This result suggests that the frequency jumps in USVs are likely to be an important feature in their classification. We conclude that our framework and the SIS measure is a feasible method that is sensitive enough to measure differences in existing classification methods and highlight the method that best captures regularities in the data. The SIS for depth 1 is computed as the KL-divergence between the 0^th^-order and 1^st^-order distribution of pairs of USVs. As the KL-divergence is a sum, we can look for the individual contribution of each combination of two syllables (of the possible 64 pairs) to the SIS, by considering the value of 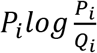. A pair for which the probability of occurring is equal to the multiplication of the two probabilities for each of the syllables to occur individually (the expected probability), will not contribute anything to the SIS. However, for some pairs the actual probability is higher or lower than the expected one. To better understand what drives the SIS, figure 4C plots these values for both iMSA and iMUPET (see Fig. S2 for all three algorithms). It is clear that in both cases the pairs that obtain the highest divergence from their expected probabilities are a repetition of the same syllable (i.e. pairs of identical syllables). Interestingly, however, for iMSA, the ‘Simple’ syllables, which in the original algorithm do not consider the duration of the syllable, show repetition only for syllable of similar duration. The pairs of Simple-long and Simple-short, occur more than expected. However, pairs of Simple-long followed by Simple-short (or vice versa) actually show up less than expected. While splitting the Simple category into two classes by the median duration, as done here, is quite arbitrary, it may indicate that the Simple class might be composed of more than one class, where the feature of syllable duration plays a significant role, and that finding these classes will reveal more of the richness of the statistical structure of USV sequences.

**FIGURE 4.**
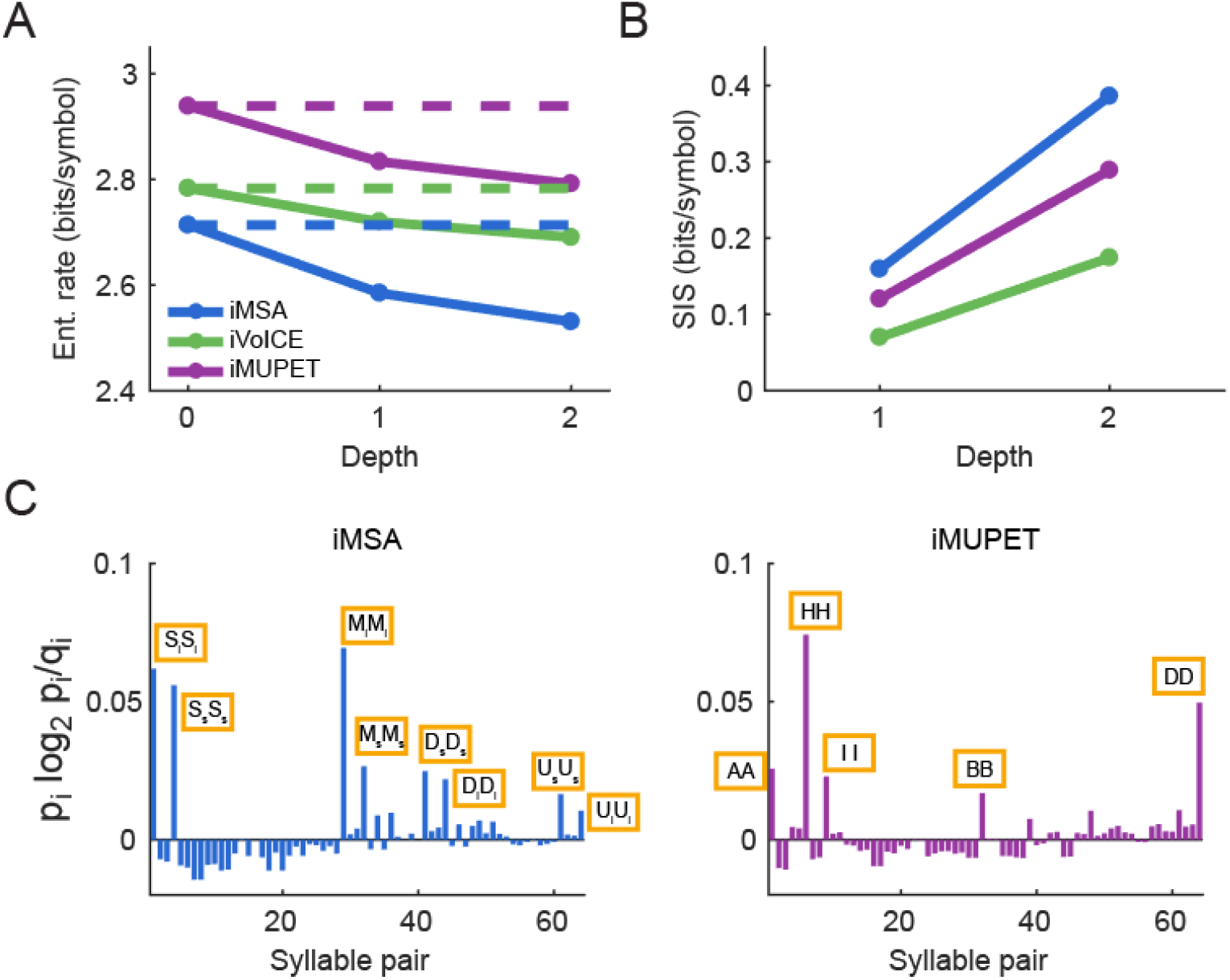
Comparison of the classification methods. **A** Entropy rate computed for the models produced by the three different methods (with 8 classes) for tree depth of order 0, 1 and 2. Solid lines represent the entropy rate for each depth; dotted lines represent the rate for zero depth. **B** The syntax information score computed for the models of depth 1 and 2 in bits/symbol. The values are calculated as the KL divergence between a given depth and the zero depth. **C** The individual contribution of each pair to the total SIS value (depth 1).

### Improving the syntax information score of current algorithms

Classification algorithms are forced by their very nature to assign a single label to each USV, even in cases where the decision is not obvious. This is especially evident for clustering-based algorithms such as VoICE and MUPET. Often the ‘clouds’ around each centroid have overlapping volumes creating some level of ambiguity. Even when using ‘soft clustering’ (Xie and Beni, 1991), where a probability to assign a label to the USV is computed for all the labels, the algorithm would eventually be forced to assign the most likely label. The higher order statistics that were showing up in our previous analysis, suggest that the sequence data may hold information that can assist classification in such case of ambiguity. This is analogues to trying to parse a note written with poor handwriting and deducing that a certain letter is likely to be a U rather than a V because it follows the letter Q. Fig. 5A illustrates such an example, where the probability assigned by an algorithm for a USV to be classified as S is larger than its probability to belong to class T. If we assign this USV the label S, this assignment, however, has also an effect on the syntax model. The next USV in the sequence will have higher probability to follow S and lower probability to follow T. This effect can be evaluated using the SIS. Doing so may reveal that assignment of T would, in fact, increase the SIS more than assignment of S. If this difference in the SIS in the two cases is large enough, we may decide that this USV should be labeled T after all, and ignore our feature-based similarity measure that is used by the classification algorithm.

**FIGURE 5.**
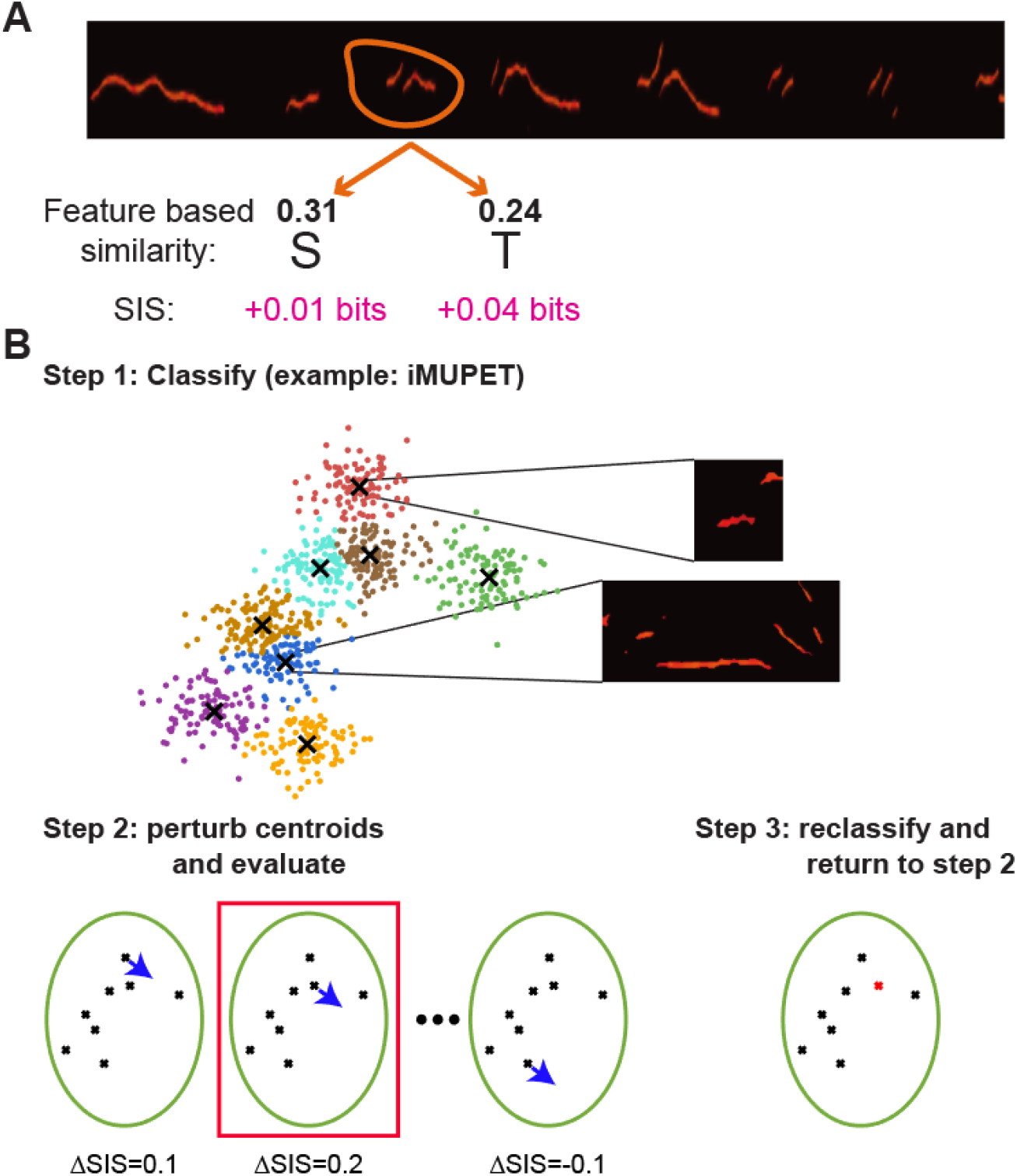
Syntax Information Maximization (SIM) algorithm. **A** Classification of a syllable to a class is based on the feature similarity of the syllable to each class. The similarity defines the probability of the syllable to belong to each class. The syntax information score (SIS) of the classification represents the amount of information that the classification provides about the next syllable in a sequence. SIM optimizes classification by using the SIS as an additional constraint. **B** Illustration of the SIM algorithm. The initial condition is the classification of another algorithm. Here we chose iMUPET as an example. iMUPET provides a set of centroids and each USV is assigned to a centroid. In step 2 a random perturbation is chosen and each of the centroids is perturbed in turn. Then, the change on the SIS is evaluated (ΔSIS). The perturbation that resulted with the largest ΔSIS is chosen (red frame). In step 3 all the USVs are reclassified based on the new set of centroids (all but the chosen centroids are the same, and the chosen centroid is replaced by its perturbed version (red dot)). Step 2 and Step 3 are repeated until convergence is achieved.

We present the Syntax Information Maximization (SIM) algorithm that considers the SIS of the classification as an optimization constraint. Given a set of centroids, the goal is to find a new set that has a larger SIS on a test set that it was not trained upon. That means that it has to consider the properties of the single syllable as well as the syntax. Here we have chosen to use iMUPET (which achieved the second-best SIS in our test, Figure 4B) and improve it. Figure 5B illustrates the process. We choose a training set of USVs (composed of half of the sequences in the database) and use iMUPET to compute centroids that represent the different classes. Next, each of the centroids is perturbed in turns, with an identical perturbation. For every perturbation, all the USVs are re-assigned to classes and the change in SIS (ΔSIS) is evaluated on the resulting syntax (still on the training set). After the ΔSIS has been evaluated for all the sets of perturbed centroids, the perturbation that resulted with the largest ΔSIS is chosen. The USVs are reclassified and the procedure is repeated.

The results of the algorithm are shown in Figure 6A. The algorithm is designed to increase the SIS in each step on the training set, and therefore it is not surprising that indeed the SIS is improving on this set. However, we also evaluated the algorithm on the test set after each step and as seen the SIS shows a similar trend on the test set that is not considered during the iteration of the algorithm. This demonstrates that the new set of centroids found by the algorithm generalizes well and captures better the syntax of the USV sequences. Note that for depth 1 the algorithm yields an SIS that is larger than that of iMSA. For depth 2 the algorithm obtains a larger SIS only on the training set, and on the test set the result is slightly lower than iMSA. Figure 6B plots 0^th^-order and 1^st^-order distributions for the SIS results of SIM. Notice that these distributions are different from those of any of the algorithms shown in figures 2B and C. Moreover, as seen in figure 6C, SIM captures pairs of syllables that have extremely unpredictable occurrence 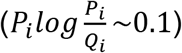. Lastly, figure 6D compares SIM to the other algorithms and shows that the new algorithm obtained improved SIS for both 1^st^-order and 2^nd^-order models over the original iMUPET algorithms and also compared to the iMSA.

**FIGURE 6.**
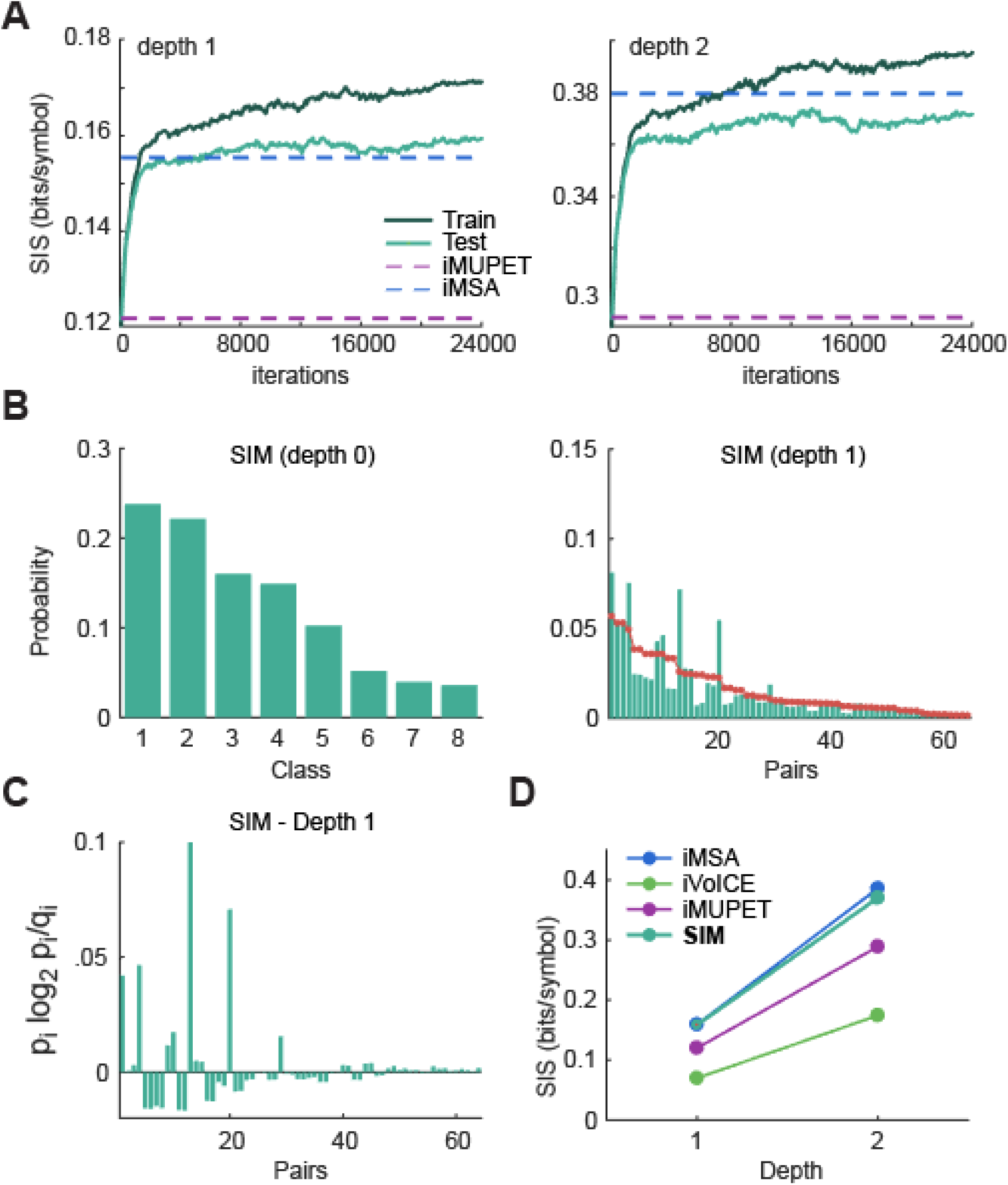
Results of the syntax information maximization (SIM). **A** Results of the syntax information maximization algorithm from figure 5 on training and test sets from with a suffix tree of depth 1. The initial condition is the classification of iMUPET (bottom dashed line). The algorithm quickly approaches the SIS of iMSA (top dashed line). Right, same for depth 2 **B**. The distribution of classes (left) and for syllable pairs (right) of the SIM algorithm. **C** The contribution of each syllable pair to the SIS of depth 1. **D** The SIS for SIM compared to the original three algorithms for depth 1 and 2.

## Discussion

The analysis of mouse ultrasonic vocalizations led to the development of many methods to classify USV syllables into discrete classes (Burkett et al., 2015; Chabout et al., 2015; Van Segbroeck et al., 2017). These proposed classification methods are based solely on the spectral representation of each single syllable. However, there is currently no ground-truth that can serve as a standard to evaluate the performance and accuracy of these methods and thus the selection of which method to use is somewhat arbitrary. In this work, we hypothesized that the syntax of USVs (i.e. the statistical properties of USVs sequences) can serve as an additional important source of information that can assist the classification process. We showed that syntax can serve as a good measure for different classification methods as well as for improving the classification process.

By subjecting the same database of USVs to different classification methods one fundamental outcome, common to all methods, became clear: the USVs are not emitted in a temporarily independent manner. Rather, the chance of a USV belonging to a certain class is affected by the classes of its preceding USVs. Given this observation, the rationale we propose here is that an essential feature of a classification algorithm is that it capture as much of that temporal dependency as possible. In other words, a good classification method provides information about the upcoming syllables based on recent ones. The more information that a classification provides, the more subtleties and motifs it discovers. However, there is a fly in the ointment; the method that obtains the ultimate predictability is one that assigns all USVs to the same class. We therefore devised a measure that penalizes such behavior, using measures from information theory. Because the classification methods do not consider temporal structure and classify each USV individually, it is reasonable to consider the entropy of the syllable distribution as a baseline predictability score, and then compare it to how knowledge of preceding syllables improves the predictability. In that way, a trivial method that assigns all USVs to the same class will have zero entropy as baseline and this cannot be improved by considering the preceding USVs.

The comparison performed here revealed that although the compared classifications differ in their classes and predictability, they all capture some meaningful features in the signal. This comparison framework is available online (link) and can be used to compare additional existing and novel methods. In our hands the best predictability score has been achieved by classification of syllables base on frequency jumps (Chabout et al., 2015), suggesting this is an important feature of USVs. Assuming there is a real natural and true classification of USVs, there might be various reasons why different classification algorithms perform differently on the proposed predictive power measurement. One option is that each algorithm concentrates on a subset of features for the classification process, while a true classification requires a wider set of features. For example, it might be that pitch jumps are an essential feature for USVs classification, but Simple upward sweeps are inherently different from downward sweeps, a feature that is not considered by MSA. A second possible reason is that the geometric projection and the metric used by the algorithm are not optimal for a true separation between USVs belonging to different classes. For example, MUPET pre-processes the frequency representation of USVs before projecting them into high-dimensional space and uses cosine-based norm to classify them. We could now use the SIS and compare the classifications using other metric measures and different pre-processing procedures over the same set of data, in order to study if this pair is the optimal one or whether a different combination might yield a yet better result.

The vocalization database that was created in this work played an important role in modeling the various classification syntaxes. Without a large amount of data, the higher order statistics of the classifications could not be calculated. This database and the parsing tool that created it are also available online and can also be used to address additional topics such as: the temporal structure of the vocalizations, sequence types, and more.

In addition to comparing classifications, the syntax was used to create an improved classification. The SIM algorithm used here (Fig. 5) demonstrated how by considering the syntax when dividing the syllable space into classes, the resulting predictability of the algorithm is higher than the original one. Clearly the algorithm we proposed is sub-optimal. The cost of each iteration, involving evaluation of the syntax model at each step is very high computationally. There are many technical steps that can be used to improve the efficiency of the algorithm. Nevertheless, the motivation here was to demonstrate that the information existing in the syntax can be used to drive the algorithm to obtain an improved representation of the classes which generalizes to data that had not been considered before (the test set). Future work that considers a more advanced approach in combining the spectral information with the syntactic information could result in classifications that have even more prediction power. The challenge with this kind of implementation is that changing the class of a syllable based on the syntax causes a change in the syntax. Therefore, a unified approach where the two sources of information are represented in the same space might produce the desired result.

Clearly, the exploration of the true number of classes in the USV classification is not over yet. In this work, we compared and improved classifications that contain eight classes, but different methods suggest different numbers of classes. To explore this, it is possible to use a similar information-theory based approach with a development of some penalty for too low (reducing entropy) or too high (reducing the classification error/over-fitting) a number of classes.

Each classification algorithm creates a syntactic model of the USVs. As we continue to develop our classification method, the model it generates becomes more detailed and precise. As this measure improves, it enables a better understanding of social disorders and treatments for them. For example, we will be able to better measure the effect of a treatment for ASD (Perets et al., 2017) or the effect of a knockout on a gene (Castellucci et al., 2016). Therefore, carrying on the integration of such analytical tools together with behavioral paradigms could result in advanced treatments for social and speech disorders.

## Materials and Methods

### Animals

For the recordings performed in our lab, we used C57BL/6 male and female mice (8–12 weeks old). All mice were group housed (3–4 per cage) and kept on a 12 h light (7 a.m.–7 p.m.)/dark cycle with ad libitum food and water.

### Ethical note

Experimental protocols were approved by the Hebrew University Animal Care and Use Committee and met guidelines of the National Institutes of Health guide for the Care and Use of Laboratory Animals (IACUC NS-16-14216-3).

### USV database

We created a database of 525 USV recording sessions of C57 male-female interactions. Most of the recordings were performed in our lab and 36 additional files were downloaded from the mouseTube (Torquet et al., 2016) (‘Female’ context recordings from the ‘Social context comparisons’ protocol (Chabout et al., 2015)). All files were in ‘wav’ format with a sampling rate of 250000 Hz, and their length ranged between 2-30 minutes. The full set of files used in this study was uploaded to mouseTube and can be found using the group label “London Lab”.

### USV parser

After testing a few parsing tools, we noticed that they were not optimized to cope with the different types of noise that existed in the USV files. These different levels of noise in the recordings were a result of: varying cage sizes, cage acoustics, locations of the recording device and noise from the freely moving mice. Therefore, we developed a USV parser (written in Python; available online) that is robust to these types of noise. Figure S3 describes the flow of the parser. The parser receives as input one or more ‘wav’ USV files and returns the start and end time of each syllable in the file.

### USV statistics

The USV parser was applied to all recordings in the database. This resulted in 461,927 syllables. Using the start and end time of each syllable, we calculated the distributions of the syllable duration and ISI. We also examined the ISI distribution for several specific mice in order to test the variability between them.

In addition, the strongest frequency in each time point was detected and the mean frequency of each syllable was stored in the database, enabling the analysis of the mean frequency distribution. An ISI of more than 160 ms represented the end of the current sequence and the beginning of a new one. We calculated the number of syllables in the different sequences and created the sequence length distribution. For calculating the second order statistics, we collected all pairs of consecutive syllables in a sequence. We ran a Pearson correlation test for both the duration and the ISI to determine if there are second-order correlations.

### Adaptation of existing algorithms

Source code for all three algorithms was available in MATLAB. We performed several adaptations to each algorithm in order to enable a fully-automated execution.

#### Mouse Song Analyzer v1.3 (Chabout et al., 2015)

The MSA algorithm includes a built-in syllable parser. In order to label the same syllables for all algorithms, we replaced the syllables detected by the MSA parser with those that were detected by our parsing algorithm (see above). We ran the MSA algorithm on all files and saw that there were files where more than 5% of the syllables were labelled as “unclassified”. For those files, we re-ran the algorithm with lower and lower thresholds (default was 0.3, decrease steps were of 0.05 and minimum value was 0.15). We selected the first threshold with an “unclassified” rate lower than 5%. If there was no such threshold, we selected the threshold with the lowest unclassified rate. Nevertheless, manual examination of the remaining “unclassified” syllables showed still a significant amount of real USVs. The shorter “unclassified” syllables were “simple” and the longer ones tended to be “multiple”. As a result, and in order to classify each one of the syllables to one of the four basic classes (simple, down, up, multiple), we gave the “unclassified” syllables one of two labels: “simple” or “multiple”. We used the median duration of all syllables in the database (35.3 ms) and set the syllables with a shorter duration as “simple” and with a longer duration as “multiple”. In total there were 50953 “unclassified” syllables of which 36683 were classified as simple (18.3% of the total simple population) and 14270 were classified as multiple (18.1% of total multiple). Next, to support an eight-class classification, we split each one of the four classes to two. This was done using the median duration of each class (simple: 27.6 ms, down: 48.1 ms, up: 50.7 ms and multiple: 96.3 ms). For example, “down” syllables that were shorter than 48.1 ms were classified as Down-short and “up” syllables longer than 50.7 ms were classified as Up-long.

#### VoICE (Burkett et al., 2015)

The VoICE algorithm is based on hierarchical clustering. Running the algorithm on all syllables in the database was not feasible because of computation constrains. Therefore, 4000 syllables from different files were selected and the algorithm was applied to them. VoICE includes a manual phase (originally used for comparison) that was skipped. The results of the automatic phase are centroids that were further used to classify all 461,927 syllables. This classification was done using the same similarity measure that was used in the other parts of the VoICE algorithm.

#### MUPET (Van Segbroeck et al., 2017)

MUPET uses a gammatone filter as part of the preprocessing. For the adapted version, we used 16 filters. As in the MSA algorithm, MUPET also contains a parsing phase. We loaded our syllable times instead of the built-in ones to maintain consistency. Like the case with VoICE, the MUPET algorithm was not able to run on the full database, therefore we applied it on 5000 syllables. Then, we used the resulting centroids and the MUPET distance measure to classify the rest of the syllables.

### Sample algorithms

The four sample algorithms used to demonstrate the quantification framework are all based on the iMSA algorithm. The first algorithm generates two labels: one class (N: no jump) that is the result of merging the two simple classes (Simple-short and Simple-long) and another class (J: jump) that is the result of merging all other six classes. Besides the original version of the MSA algorithm, the second four-class algorithm contains both Simple-short and Simple-long classes, another class that contains all four Down/Up-short/long classes and a final class containing both the multiple classes (short and long). The final classification is composed of five classes which are: Simple-short, Simple-long, Down-merged, Up-merged and Multiple-merged. This diversity allows examining our framework for classifications with different numbers of classes and different distributions of syllables.

### From classification to Suffix tree

We define our Suffix tree as a full tree where each leaf is associated with one suffix. The suffix is composed of the edge labels on the path from the root to the leaf. Each leaf holds the probability of the suffix associated with it to appear (denoted as *μ*_*i*_ for leaf *i*). In addition, it holds the probabilities for each one of the syllables to appear after that suffix (denoted as *P*_*ij*_ for leaf *i* and syllable *j*). The number of leaves in the suffix tree is *K*^*d*^ where *K* is the number of classes and *d* is the depth of the tree. Therefore, for a classification with four classes: RSTU, and a tree with depth 2, the leaves will represent the following 16 suffixes: RR, RS, RT, RU, SR, SS, ST, …, US, UT, UU. In this example, each leaf holds a discrete probability distribution with four values.

To create such a tree for a given classification algorithm, we apply the algorithm on all syllables and divide them into sequences. The sequences are scanned one-by-one and for each suffix in the sequence, the value of the corresponding leaf is increased by 1. In addition, the value of the next syllable that follows the suffix is increased by 1 as well. At the end of this process, the counts of all leaves are converted into probabilities. The suffix probabilities are calculated by dividing the suffix count with the total number of syllables. The following syllable probabilities are calculated by dividing each count by the leaf sum.

### Entropy rate calculation

#### Theorem 2.2.4 (Cover and Thomas, 1991)

Let {*X*_*i*_} be a stationary Markov chain with stationary distribution *μ* and a transition matrix P. Let *X*_1_ ~ *μ*. Then the entropy rate is:

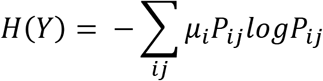

In our case, the stationary distribution is stored in the leaves. Therefore, the probability of a suffix represented by leaf *i* is denoted as *μ*_*i*_. The transition matrix is distributed between the leaves such that each leaf holds one row of the matrix with *j* columns corresponding to the *j* options for the following syllable.

### KL divergence calculation

#### Definition (Cover and Thomas, 1991)

The relative entropy or Kullback–Leibler distance between two probability mass functions p(x) and q(x) is defined as:

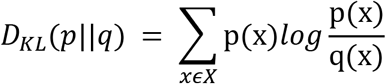

We use the KL distance measure to calculate the distance between the probability function of the higher-order model and the zero-order model. The leaves of the suffix tree hold the probabilities of the higher-order model (p(x)), while the values of q(x) are calculated as the expected probability from the zero-order model. The expected probability is calculated by multiplying the zero-order probabilities of all syllables in the suffix. Given the example of four classes: RSTU, the expected probability for q(S|TU) will be: q(S|TU) = *p*_0_(*S*) × *p*_0_(*T*) × *p*_0_(*U*), where *p*_0_ represents the zero-order probability.

### Syntax information maximization algorithm

For a given classification, the SIM divides the sequences into two groups: training and test sets, with the same number of sequences in both sets. Every syllable in the database goes through the MUPET pre-processing procedure with 16 filters. This converts all the syllables into vectors with a length of 2016. The initial SIS is calculated for the training set, as well as the centroid of each class. The centroid is calculated as the mean of all syllables in the class. Then, the iterative process starts. In each iteration, a random vector *V* is created by generating values from a uniform distribution ranging between 0.9 and 1.1. This vector is used to perturb each centroid, one at a time. The perturbation is done as a dot product between the vector representing the centroid and *V*. For each perturbation, all syllables in the training set are re-classified and the SIS is calculated for the new classification. After perturbing all centroids, there is an SIS value corresponding to each perturbation. The maximum value is selected and compared to the value before perturbation. If it is higher, the perturbation that achieved that value is applied and stored and a new random vector V is generated. If the maximum value is lower, then a “failure counter” is increased and no perturbation is done. Once the “failure counter” reaches the value of 5, the perturbation with the maximum SIS is applied, no matter if it is larger or smaller than the pre-perturbation SIS. Anytime a perturbation is applied, the “failure counter” is reset to 0.

This iterative process is repeated until the SIS of the training set converges. Once convergence is achieved, the SIM “replays” the same perturbation chain on the test set and the SIS in each step is calculated. The progress of the SIS on the training set and on the test set is then plotted.

---

**SUPPLEMENTARY FIGURE S1.**
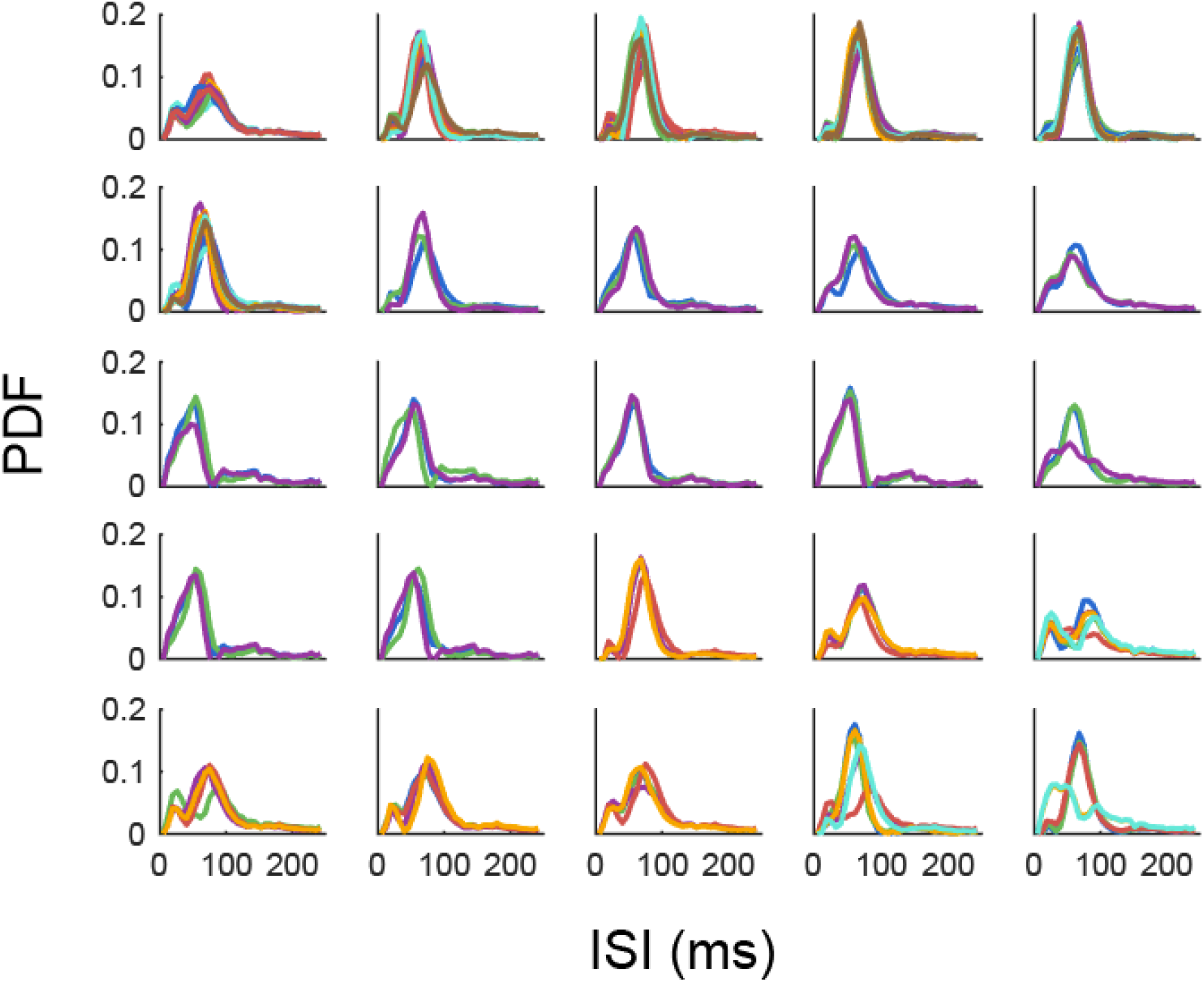
ISI distributions of individual mice. A matrix of 5×5 ISI distributions, where each cell corresponds to an individual mouse and each colored line represents a single session. Some of the mice have stereotypic bi-modal distributions (peaks around 20 ms and 70 ms) while others have a single peak around 60 ms.

**SUPPLEMENTARY FIGURE S2.**
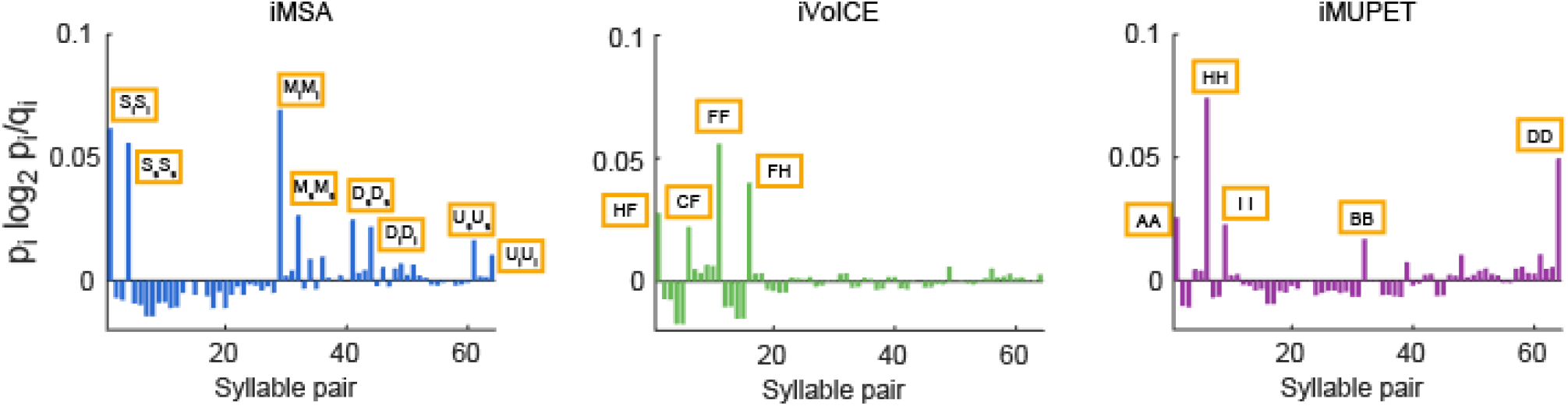
SIS contribution of individual pairs. The contribution of each pair to the total SIS value (depth 1).

**SUPPLEMENTARY FIGURE S3.**
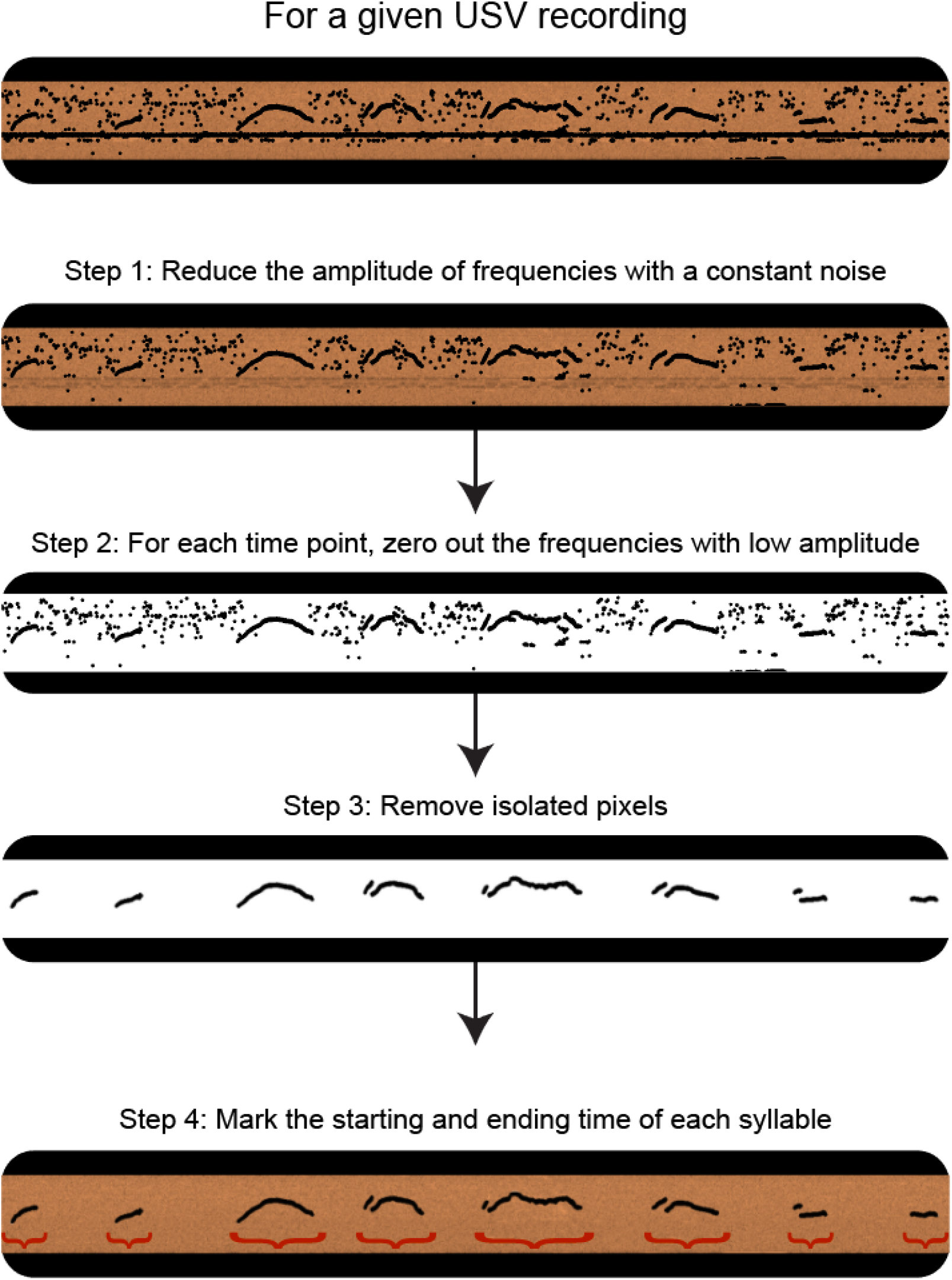
An illustration of the main phases of the parsing algorithm. The algorithm converts the USV recording to a spectrogram and applies three main stages of noise removal (steps 1-3). Then, the algorithm scans the clean spectrogram, detects the starting and ending time of each syllable and returns them as a list (step 4). Step 1: Some of the recordings contain one or more frequencies with a constant noise. The first step of the algorithm searches for this type of noise and reduces the amplitude of the corresponding frequencies. Step 2: For each time point, frequencies with low amplitude are set to zero. Step 3: Pixels that the sum of amplitudes of their neighbors is low are set to zero. Step 4: The starting and ending point of each syllable is detected.

## References

Arriaga G, Zhou EP, Jarvis ED (2012) Of Mice, Birds, and Men: The Mouse Ultrasonic Song System Has Some Features Similar to Humans and Song-Learning Birds. PLoS One 7:e46610 Available at: https://doi.org/10.1371/journal.pone.0046610.

Brainard MS, Doupe AJ (2002) About Learning. 417:351–358.

Burkett ZD, Day NF, Peñagarikano O, Geschwind DH, White SA (2015) VoICE: A semi-automated pipeline for standardizing vocal analysis across models. Sci Rep.

Castellucci GA, McGinley MJ, McCormick DA (2016) Knockout of Foxp2 disrupts vocal development in mice. Sci Rep.

Chabout J, Sarkar A, Dunson DB, Jarvis ED (2015) Male mice song syntax depends on social contexts and influences female preferences. Front Behav Neurosci.

Chabout J, Sarkar A, Patel SR, Radden T, Dunson DB, Fisher SE, Jarvis ED (2016) A Foxp2 mutation implicated in human speech deficits alters sequencing of ultrasonic vocalizations in adult male mice. Front Behav Neurosci 10:197.

Cover TM, Thomas JA (2005) Elements of Information Theory.

Doupe AJ, Kuhl PK (1999) BIRDSONG AND HUMAN SPEECH: Common Themes and Mechanisms. Annu Rev Neurosci 22:567–631 Available at: http://www.annualreviews.org/doi/10.1146/annurev.neuro.22.1.567.

Egnor SER, Branson K (2016) Computational Analysis of Behavior. Annu Rev Neurosci.

Fischer J, Hammerschmidt K (2011) Ultrasonic vocalizations in mouse models for speech and socio-cognitive disorders: Insights into the evolution of vocal communication. Genes, Brain Behav 10:17–27.

Fisher SE, Vernes SC (2015) Genetics and the Language Sciences. Annu Rev Linguist 1:289–310.

Holy TE, Guo Z (2005) Ultrasonic songs of male mice. PLoS Biol.

Hosino T, Okanoya K (2000) Lesion of a higher-order song nucleus disrupts phrase level complexity in Bengalese finches. Neuroreport 11:2091–2095 Available at: http://www.ncbi.nlm.nih.gov/pubmed/10923650.

Jarvis ED (2004) Learned birdsong and the neurobiology of human language. Ann N Y Acad Sci 1016:749–777.

Kullback S, Leibler RA (1951) On Information and Sufficiency. Ann Math Stat.

MacQueen J (1967) Some methods for classification and analysis of multivariate observations. In: Proceedings of the fifth Berkeley symposium on mathematical statistics and probability, pp 281–297. Oakland, CA, USA.

Okanoya K (2004) Song Syntax in Bengalese Finches: Proximate and Ultimate Analyses. Adv Study Behav 34:297–346.

Okubo TS, Mackevicius EL, Payne HL, Lynch GF, Fee MS (2015) Growth and splitting of neural sequences in songbird vocal development. Nature 528:352–357 Available at: http://dx.doi.org/10.1038/nature15741.

Perets N, Segal-Gavish H, Gothelf Y, Barzilay R, Barhum Y, Abramov N, Hertz S, Morozov D, London M, Offen D (2017) Long term beneficial effect of neurotrophic factors-secreting mesenchymal stem cells transplantation in the BTBR mouse model of autism. Behav Brain Res 331:254–260.

Portfors C V., Perkel DJ (2014) The role of ultrasonic vocalizations in mouse communication. Curr Opin Neurobiol.

Portfors C V (2007) Types and functions of ultrasonic vocalizations in laboratory rats and mice. J Am Assoc Lab Anim Sci.

Price PH (1979) Developmental determinants of structure in zebra finch song. J Comp Physiol Psychol 93:260–277.

Sales G (1972) Ultrasound and mating behavior in rodents with some observations on other behavioral situations.

Scattoni ML, Gandhy SU, Ricceri L, Crawley JN (2008) Unusual repertoire of vocalizations in the BTBR T+tf/J mouse model of autism. PLoS One 3:48–52.

Seagraves KM, Arthur BJ, Egnor SER (2016) Evidence for an audience effect in mice: male social partners alter the male vocal response to female cues. J Exp Biol.

Sewell GD (1968) Ultrasound in rodents. Nature 217:682–683.

Sugimoto H, Okabe S, Kato M, Koshida N, Shiroishi T, Mogi K, Kikusui T, Koide T (2011) A role for strain differences in waveforms of ultrasonic vocalizations during Male-Female interaction. PLoS One 6.

Torquet N, de Chaumont F, Faure P, Bourgeron T, Ey E (2016) mouseTube – a database to collaboratively unravel mouse ultrasonic communication. F1000Research.

Van Segbroeck M, Knoll AT, Levitt P, Narayanan S (2017) MUPET—Mouse Ultrasonic Profile ExTraction: A Signal Processing Tool for Rapid and Unsupervised Analysis of Ultrasonic Vocalizations. Neuron 94:465–485.e5 Available at: http://dx.doi.org/10.1016/j.neuron.2017.04.005.

Williams H, Staples K (1992) Syllable chunking in zebra finch (Taeniopygia guttata) song. J Comp Psychol 106:278–286 Available at: http://www.ncbi.nlm.nih.gov/pubmed/1395497.

Xie XL, Beni G (1991) A validity measure for fuzzy clustering. IEEE Trans Pattern Anal Mach Intell.

Yang M, Mahrt EJ, Lewis F, Foley G, Portmann T, Dolmetsch RE, Portfors C V., Crawley JN (2015) 16p11.2 deletion syndrome mice display sensory and ultrasonic vocalization deficits during social interactions. Autism Res.

